# Comparison of the effectiveness of different normalization methods for metagenomic cross-study phenotype prediction under heterogeneity

**DOI:** 10.1101/2023.10.15.562417

**Authors:** Beibei Wang, Fengzhu Sun, Yihui Luan

## Abstract

The human microbiome, comprising microorganisms residing within and on the human body, plays a crucial role in various physiological processes and has been linked to numerous diseases. To analyze microbiome data, it is essential to account for inherent heterogeneity and variability across samples. Normalization methods have been proposed to mitigate these variations and enhance comparability. However, the performance of these methods in predicting binary phenotypes remains understudied. This study systematically evaluates different normalization methods in microbiome data analysis and their impact on disease prediction. Our findings highlight the strengths and limitations of scaling, compositional data analysis, transformation, and batch correction methods. Scaling methods like TMM and RLE show consistent performance, while compositional data analysis methods exhibit mixed results. Transformation methods, such as Blom and NPN, demonstrate promise in capturing complex associations. Batch correction methods, including BMC and Limma, consistently outperform other approaches. However, the influence of normalization methods is constrained by population effects, disease effects, and batch effects. These results provide insights for selecting appropriate normalization approaches in microbiome research, improving predictive models, and advancing personalized medicine. Future research should explore larger and more diverse datasets and develop tailored normalization strategies for microbiome data analysis.

## Introduction

The human microbiome is a complex ecosystem of microorganisms that exist in symbiosis with the human body [1]. Extensive research has established that the human microbiome plays crucial roles in numerous physiological processes, including digestion, metabolism, immune system modulation, and even cognitive functions. Disruptions in the delicate microbial balance, known as dysbiosis, have been linked to a wide range of health conditions, including obesity [2, 3], diabetes [4], inflammatory bowel disease [5, 6], allergies [7], and several types of cancer [8, 9].

The advent of high-throughput sequencing technologies has revolutionized the field of microbiome research, enabling comprehensive profiling of microbial communities and providing insights into their roles in different physiological processes and disease states [10]. However, the analysis of microbiome data poses significant challenges due to inherent heterogeneity and variability across samples. Sources of variation can stem from technical differences in sequencing protocols [11], variations in sample collection [12] and processing methods [13], as well as biological diversity among individuals and populations. To extract meaningful insights from microbiome data, it is crucial to account for and mitigate these sources of variation.

Normalization methods have emerged as vital tools in addressing the heterogeneity and biases present in microbiome data. These methods aim to remove technical and biological biases, standardize data across samples, and enhance comparability between datasets. Various normalization approaches have been proposed, ranging from simple scaling methods to more advanced statistical techniques. Comparisons of normalization methods have been performed in the context of data distributions [14, 15] and differential analysis[16, 17, 18, 19, 20]. Genotype-to-phenotype mapping is an essential problem in the current genomic era. However, the impact of normalization methods on predictions mainly focused on DNA microarray data and RNA-Seq data. Zwiener et al. [21] found rank-based transformations performed well in all scenarios in real RNA-Seq datasets. Franks et al. [22] proposed feature-wise quantile normalization (FSQN) and found FSQN successfully removes platform-based bias from RNA-Seq data, regardless of feature scaling or machine learning algorithm. Given the central role of normalization in microbiome data analysis and the lack of current methods comparison for microbiome data, there is a need to systematically evaluate their performance, particularly in the context of disease prediction.

In this paper, we provide a review of existing normalization methods and present a comprehensive evaluation of various normalization methods in predicting binary phenotypes using microbiome data. We examine the performance of scaling methods, compositional data analysis methods, transformation methods, and batch correction methods across simulated datasets and real datasets. Our analysis includes an assessment of prediction accuracy using metrics such as the area under the receiver operating characteristic curve (AUC) and the rank ordering of different methods.

By comparing and contrasting the performance of normalization methods across different datasets and phenotypic outcomes, we aim to provide insights into the strengths and limitations of each approach. This research will assist researchers and practitioners in selecting appropriate normalization methods for microbiome data analysis, thereby enhancing the robustness and reliability of predictive models in microbiome research.

## Materials and methods

### Real metagenomic dataests

As the first application example, we analyzed shotgun sequencing data from patients with colorectal cancer (CRC) obtained from the R package curatedMetagenomicData v3.8.0 [23]. The taxonomic profiles for each dataset were determined using MetaPhlAn3 [24], which ensures consistency in downstream analysis. A total of nine CRC datasets are available [25, 26, 27, 8, 28, 29, 30, 9, 31]. We excluded studies with sample sizes of less than 30 for either cases or controls, resulting in eight accessible CRC datasets for our analysis. A detailed summary outlining the distinctive characteristics of these eight CRC datasets can be found in Table 1.

**Table 1:** Characteristics of CRC datasets, with different DNA extraction kits (DNA-Exk) and sequencing platforms (Seq-Plat).

| Dataset | Country | No. of control | No. of CRC | DNA-Exk | Seq-Plat | Reference |
| --- | --- | --- | --- | --- | --- | --- |
| Feng | Austria | 61 | 46 | MoBio | IlluminaHiSeq | [25] |
| Gupta | Indian | 30 | 30 | Qiagen | IlluminaNextSeq | [68, 26] |
| Thomas | Italy | 52 | 61 | Qiagen | IlluminaHiSeq | [8] |
| Vogtmann | United States of America | 52 | 52 | Gnome | IlluminaHiSeq | [28] |
| Wirbel | Germany | 65 | 60 | Gnome | IlluminaHiSeq | [29] |
| Yachida | Japan | 251 | 258 | NA | IlluminaHiSeq | [30] |
| Yu | China | 53 | 75 | Qiagen | IlluminaHiSeq | [9] |
| Zeller | France | 61 | 53 | Gnome | IlluminaHiSeq | [31] |

As the second application example, we analyzed shotgun sequencing data from patients with inflammatory bowel disease (IBD) from the R package curatedMetagenomicData v3.8.0 [23]. There are 6 available IBD datasets in curatedMetagenomicData [32, 5, 33, 34, 35, 6]. Similarly to the CRC datasets, we excluded studies with sample sizes less than 30 for either cases or controls from the analysis. A summary of the characteristics of the IBD datasets can be found in Supplementary Table S1.

### Statistical analysis

We calculated the microbial relative abundance for each sample and used the Bray-Curtis distance [36] to compare the dissimilarities between samples. This distance was computed using the function *vegdist()* from R package *vegan* [37]. To visualize the clustering of samples effectively, we performed principal coordinate analysis (PCoA) through the *pcoa()* function from R package *ape* [38]. To assess the variance attributable to datasets, we conducted the permutational multivariate analysis of variance (PERMANOVA) [39] with *adonis()* function in R package *vegan* [37].

### Normalization methods

A number of normalization methods could be applied to microbiome data for data analyses. For the purpose of predicting the unknown disease status of samples, we try to transform or normalize our data to satisfy the assumption that training and testing data are drawn from the same distribution. Seven scaling methods, one compositional data analysis method, eight transformation methods, and six batch correction methods were compared in this analysis. Our study is also the largest comparison in terms of prediction up to date according to our best knowledge.

A taxa count table of a metagenomic dataset can be organized as shown in Table 2. Assume we have a dataset consisting of *n* samples and *m* features. Denote *c*_*ij*_ as the count for taxon *i* in sample *j*. With this notation, the steps and formula of normalization methods can be briefly introduced as follows.

**Table 2:**
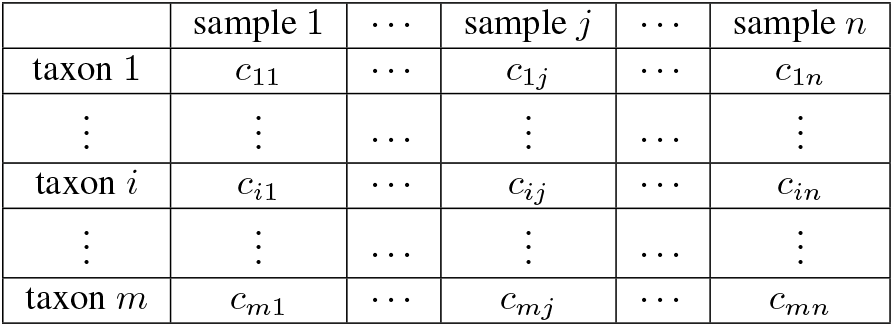
Format of a count table of metagenomic dataset, where *c*_*ij*_ is the number of reads belonging to taxon *i* and sample *j*.

### Scaling methods

A commonly used method for normalizing microbiome data is scaling. Its basic idea is to divide counts in the taxa count table by a scaling factor or normalization factor to remove biases resulting from sequencing technology:

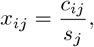

where *x*_*ij*_ is the normalized abundance for taxon *i* in sample *j, s*_*j*_ is the scaling/normalization factor for sample *j*. We investigated seven popular scaling methods (Table 3) in our analysis, including TSS, UQ, MED, CSS in metagenomeSeq, TMM in edgeR, RLE in DESeq2, and GMPR in GUniFrac.

**Table 3:**
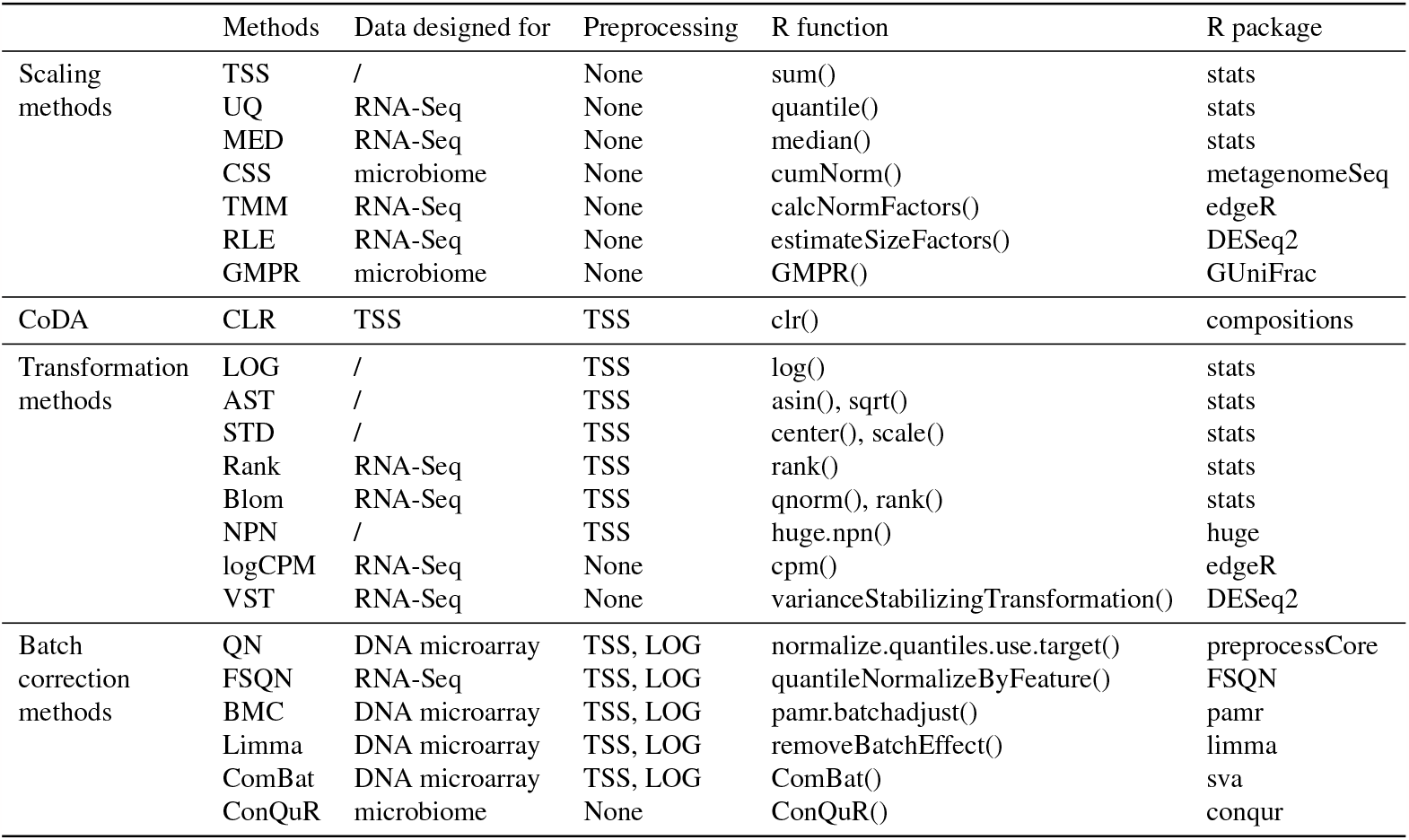
Summary of normalization methods, including seven scaling methods, one compositional data analysis (CoDA) method, eight transformation methods, and six batch correction methods.

### Total Sum Scaling (TSS) [14]

Counts are divided by the total number of reads in that sample.

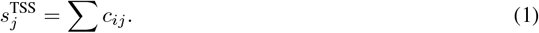

### Upper Quartile (UQ) [14, 40]

Similar to TSS, it scales each sample by the upper quartile of counts different from 0 in that sample.

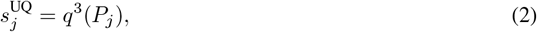

where *q*^3^(*·*) is the function of estimating upper quartile, and *P*_*j*_ = {*c*_*ij*_|*c*_*ij*_ *>* 0, *i* = 1, …, *n}* represents a set of counts different from 0 in sample *j*.

### Median (MED) [14]

Also similar to TSS, the total number of reads is replaced by the median counts different from 0 in the computation of the scaling factor.

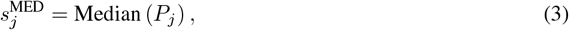

where Median (·) is the function of estimating median, and *P*_*j*_ = { *c*_*ij*_|*c*_*ij*_ *>* 0, *i* = 1, …, *n}* represents a set of counts different from 0 in sample *j*.

### Cumulative Sum Scaling (CSS) [41]

CSS modified TSS for microbiome data in a sample-specific manner. It selects the scaling factor as the cumulative sum of counts, up to a percentile 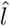 determined by the data:

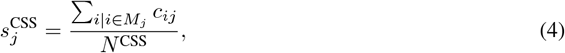

where 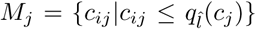 denotes the taxa included in the cumulative summation for sample *j*, and *N* ^CSS^ is an appropriately chosen normalization constant. This scaling method is implemented by calling the *cumNorm()* function in the R package *metagenomeSeq* [41].

### Trimmed Mean of M-values (TMM) [42]

TMM is a popular normalization method for RNA-Seq data with the assumption that most genes are not differentially expressed. It selects a reference sample first and views the others as test samples. If not specified, the sample with count-per-million upper quantile closest to the mean upper quantile is set as the reference. The scale factor between the test sample and the reference sample is estimated by the ratio of two observed relative abundance for a taxon *i*. The log2 of the ratio is called M value, 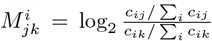, and the log2 of the geometric mean of the observed relative abundance is called A value, 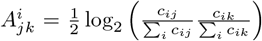. By default, it trims the *M* values by 30% and the *A* values by 5%. Then the weighted sum of *M* values can be used to calculate the scale factor of sample *j* to sample *k*:

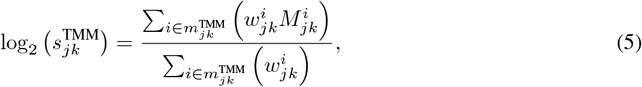

where 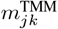 is the remaining taxa after the trimming step, and weight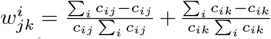.This scaling method is implemented using *calcNormFactors()* function in the *edgeR* [43] Bioconductor package.

### Relative log expression (RLE) [44]

RLE is another widely used method for RNA-Seq data and relies on the same assumption that there is a large invariant part in the count data. It first calculates the geometric mean of the counts to a gene from all the samples and then computes the ratio of a raw count over the geometric mean to the same gene. The scale factor of a sample is obtained as the median of the ratios for the sample:

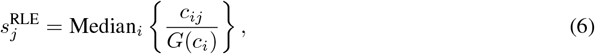

where 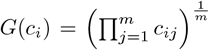 is the geometric mean of gene *i*. By setting the *type=“poscounts”* of *estimateSizeFactors()* function in the *DESeq2* [45] Bioconductor package, a modified geometric mean is computed. This calculation takes the n-th root of the product of the non-zero counts to deal with zeros in microbiome data.

### Geometric mean of pairwise ratios (GMPR) [46]

GMPR extends the idea of RLE normalization by reversing the order of computing geometric and median to overcome the zero inflation problem in microbiome data. The scale factor for a given sample *j* using reference sample *k* is calculated as

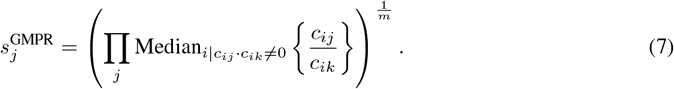

This scaling method is implemented using *GMPR()* function in the *GUniFrac* [47] package.

### Compositional data analysis (CoDA) methods

Gloor et. al. [48] pointed out that microbiome datasets generated by high-throughput sequencing are compositional because they have an arbitrary total imposed by the instrument. Thus several methods were proposed to eliminate the effect of sampling fraction by converting the abundances to log ratios within each sample. These commonly used methods in compositional data analysis include additive log-ratio transformation (ALR)[49], centered log-ratio transformation (CLR)[49], and isometric log-ratio transformation (ILR)[49]. ALR and ILR convert *n* dimensional gene vector to *n −* 1 dimensional data in the Euclidean space, with the challenge of choosing a reference gene. Due to the large number of genes and the resulting computing problem, we only considered CLR in our analysis.

### Centered Log-Ratio (CLR) [49]

CLR transformation is a compositional data transformation that takes the log-ratio of counts and their geometric means. This is done within each sample based on relative abundances. This can be written in mathematical form as:

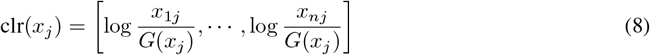

where *x*_*ij*_ is the relative abundance of gene *i, i* = 1, …, *n* in sample *j, j* = 1, …, *m*, 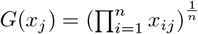 is the geometric mean of sample *j* with a pseudo count 0.65 times minimum non-zero abundance added to 0 values [50]. This transformation is implemented using *clr()* function in R package *compositions* [51].

### Transformation methods

Microbiome data have problematic properties such as skewed distribution, unequal variances for the individual taxon, and extreme values. We propose to transform microbiome data before fitting the prediction model to handle either one, two, or all of these problems. Let *c*_*ij*_ and *x*_*ij*_ be the count and relative abundance of gene *i, i* = 1, …, *n* in sample *j, j* = 1, …, *m*, respectively. Table 3 gives a summary of transformation methods considered in this study, including LOG, AST, STD, Rank, Blom, NPN in huge, logCPM in edgeR, and VST in DESeq2.

### LOG

Log transformation is often used for taxa with skewed distribution so that the transformed abundances are more or less normally distributed [21]. A pseudo count 0.65 times the minimum non-zero abundance is added to the zero values before log transformation to avoid infinite values [50].

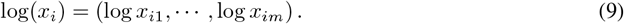

### Arcsine square-root (AST)

AST transformed data have less extreme values compared to the untransformed data and are more or less normally distributed. It is defined as

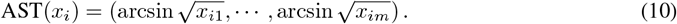

### Standardization (STD) [21]

STD is the default implementation in many regression analyses to reduce the variations of features (taxa in our analysis):

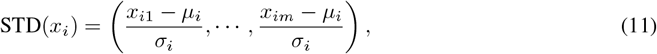

where *μ*_*i*_ and *σ*_*i*_ is the mean and standard deviation of gene *i* separately.

### Rank [21]

Rank transformation is a simple and popular method used in non-parametric statistics. The rank-transformed features are uniformly distributed from zero to the sample size *m*. A small noise term *ϵ*_*ij*_ *∼ N* (0, 10^*−*10^) is added before data transformation to handle the ties of zero counts.

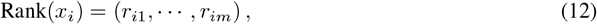

where *r*_*ij*_, *j* = 1, …, *m* is the corresponding rank for relative abundance *x*_*ij*_, *j* = 1, …, *m* in gene *i*.

### Blom [52, 21]

Blom transformation is based on rank transformation. The uniformly distributed ranks are further transformed into a standard normal distribution:

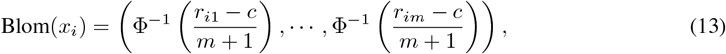

where 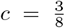 is a constant,Φ^*−*1^(*·*) denotes the quantile function of normal distribution, and *r*_*ij*_, *j* =1, …, *m* is the corresponding rank for relative abundance *x*_*ij*_, *j* = 1, …, *m* in gene *i*.

### Non-paranormal (NPN) [53]

NPN transformation is designed to be used as part of an improved graphical lasso that first transforms variables to univariate smooth functions that estimate a Gaussian copula. The transformation can also be used alone for analysis. Let Φ denote the Gaussian cumulative distribution function, then we can estimate the transformed data using

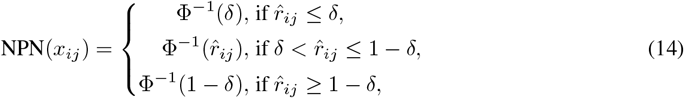

where 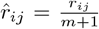, and 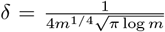 . This transformation is implemented using *huge*.*npn()* function in R package *huge* [54].

### Log counts per million (logCPM)

logCPM refers to the log counts per million, which is a useful descriptive measure for the expression level of a gene for RNA-Seq data. We applied it to the microbiome data. A pseudo count 0.65 times the minimum non-zero abundance is added to the zero values before log transformation.

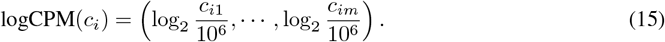

This transformation method is implemented using *cpm()* function in the *edgeR* [43] Bioconductor package.

### Variance Stabilizing Transformation (VST) [44]

VST models the relationship between mean *μ*_*i*_ and variance 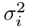 for each gene *i*:

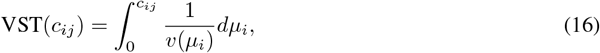

where 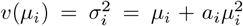, with 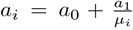being a dispersion parameter and *a*_0_ and *a*_1_ are estimated in a generalized linear model. A pseudo count 1 was added to zero values. This transformation is implemented using *varianceStabilizingTransformation()* function in the *DESeq2* [45] Bioconductor package.

### Batch correction methods

Batch effects in many genomic technologies result from various specimen processing. And they often cannot be fully addressed by normalization methods alone. Many methods have been proposed to remove batch effects. Here we studied six commonly used approaches, including QN in *preprocessCore*, FSQN in *FSQN*, BMC in *pamr*, limma in *limma*, ComBat in *sva*, and ConQuR in *conqur* (Table 3).

### Quantile normalization (QN) [55]

QN is initially developed for use with DNA microarrays, but has since been expanded to accommodate a wide range of data types, including microbiome data. Given a reference distribution, QN essentially replaces each value in a target distribution with the corresponding value from a reference distribution, based on identical rank order. In cases where the reference distribution encompasses multiple samples, the reference distribution should be first quantile normalized across all samples [56]. In our analysis, we designated the training data as the reference distribution. We applied QN to log-transformed relative abundances, substituting zeros with a pseudo count that was calculated as 0.65 times the minimum non-zero abundance across the entire abundance table. The reference distribution is obtained using function *normalize*.*quantiles*.*determine*.*target()* in R package *preprocessCore* [57]. And the batch effects are removed using function *normalize*.*quantiles*.*use*.*target()* in R package *preprocessCore* [57].

### Feature specific quantile normalization (FSQN) [22]

FSQN is similar to QN, except for quantile normalizing the genes rather than samples. The reference distribution is the gene in the training set and the target distribution is the gene in the testing set. It is applied to log-transformed relative abundance data, with zeros replaced with pseudo count 0.65 times the minimum non-zero abundance across the entire abundance table, using function *quantileNormalizeByFeature()* in R package *FSQN* [22].

### Batch mean centering (BMC) [58]

BMC centers the data batch by batch. The mean abundance per gene for a given dataset is subtracted from the individual gene abundance. It is applied to log-transformed relative abundance data, with zeros replaced with pseudo count 0.65 times the minimum non-zero abundance across the entire abundance table, using *pamr*.*batchadjust()* function from *pamr* R package [59].

### Linear models for microarray data (Limma) [60]

Limma fits a linear model to remove the batch effects. We first calculate the relative abundances and apply a log2 transformation to them. A pseudo count 0.65 times the minimum non-zero abundance across the entire abundance table was added to zeros to avoid infinite values for log transformation. The *removeBatchEffect()* function in R package *limma* [60] is then used to correct for batch effects, taking the log2 relative abundance data and batch information as inputs.

### ComBat [61]

ComBat uses an empirical Bayes framework to estimate and remove the batch effects while preserving the biological variation of interest. Similar to Limma, the relative abundance of microbiome data (zero replaced with pseudo count 0.65 times the minimum none-zero abundance across the entire abundance table) was log-transformed prior to batch correction. This correction method is implemented using the function *ComBat()* in R package *sva* [62].

### Conditional quantile regression (ConQuR) [63]

ConQuR conducts batch effects removal from a count table by conditional quantile regression. This batch correction method is implemented using function ConQuR in the R package ConQuR [63].

### The random forest classifiers

In both the CRC and the IBD datasets, we aimed to predict whether a sample originated from a case subject (CRC/IBD) or a control subject.

The training and testing datasets underwent normalization to minimize heterogeneities both within and across datasets. For scaling methods that select references, such as TMM and RLE, and transformation methods that make prediction covariates (taxa) drawn from the same distribution, such as STD, Rank, Blom, NPN, and VST, we performed normalization on the training data and then performed addon normalization of the test data onto the training data. This approach ensures that the normalization of the training data remains independent of the testing data. [64].

We performed prediction of disease status using random forest, which has been shown to outperform other learning tools for most microbiome data[65]. The random forest models were implemented using function *train()* in R package *caret* [66] with 1,000 decision trees, and the number of variables at each decision tree was tuned using grid search by 10-fold cross-validation. The area under the ROC curve (AUROC) was used as a criterion for measuring the prediction accuracy. AUC values were computed using the *roc()* function in R package pROC [67].

### Simulation studies

A successful predictive model is transferable across datasets. To evaluate the impact of various normalization methods on binary phenotype prediction, we conducted simulations by creating two case-control populations, normalizing them using various methods, building prediction models with random forest on one simulated population, and testing them on the other in 3 different scenarios. The prediction accuracy, measured by AUC values, was evaluated for each of the 100 simulation runs in different scenarios.

### Scenario 1: Different background distributions of taxa in populations

In the first scenario, we assumed that the heterogeneities between populations were due to variations in the background distributions of taxa, such as ethnicity or diet. McMurdie and Holmes [16] presented a way to simulate samples from different populations (Simulation A) and samples with case-control status (Simulation B) separately in such a scenario. In our simulations, we integrated these strategies and introduced certain modifications.

Our methodology began by determining the underlying taxon abundance levels for the training and testing populations. From Figure 1, the two least overlapping datasets, Gupta [68, 26] and Feng [25], were chosen to be the template of training and testing sets, respectively. More specifically, 30 control samples and 183 species of the Gupta dataset were included for simulating the dataset for training, and 61 healthy samples and 468 species of the Feng dataset were included for simulating the dataset for testing. For each dataset, we had a count table with rows for taxa and columns for samples. Sum the rows to get the original vectors representing the underlying taxa abundance in different populations, denoted as *p*_*k*_, *k* = 1, 2, respectively.

**Figure 1:**
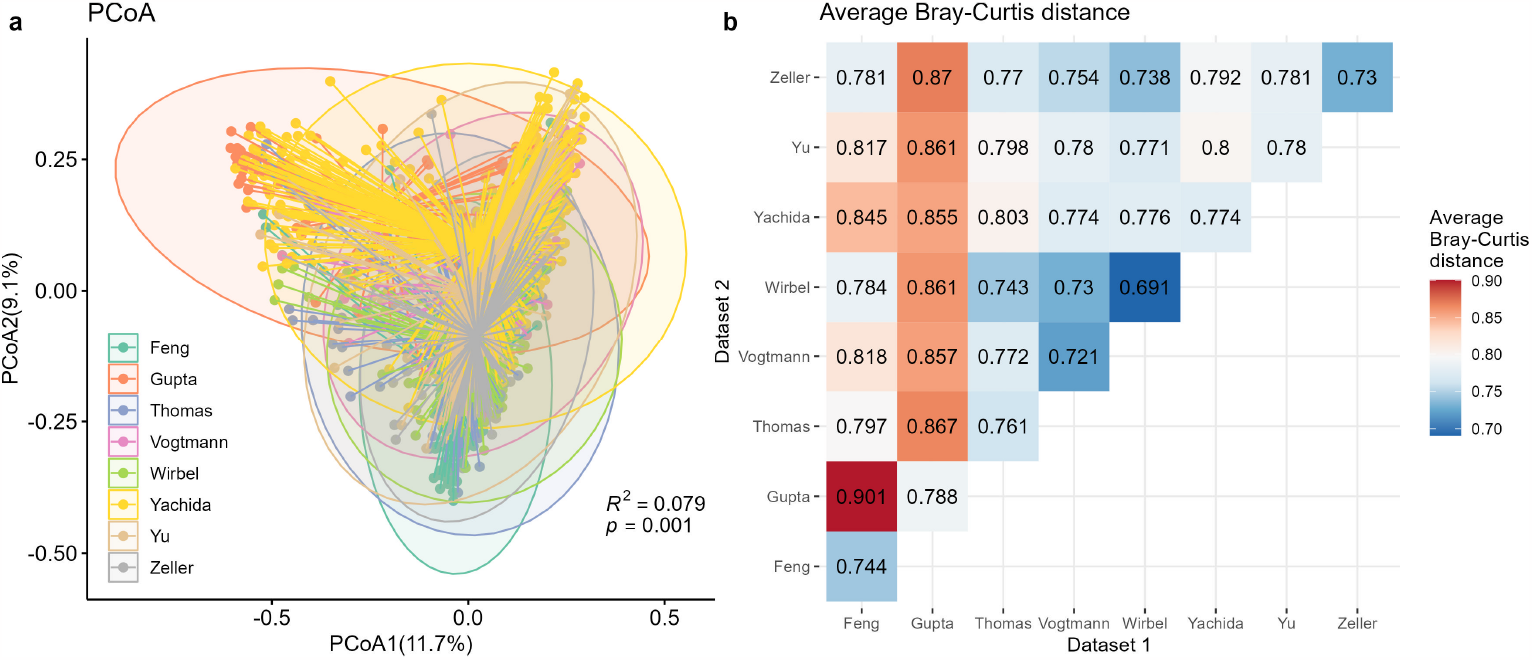
Different CRC populations had different background distribution patterns. (**a**) PCoA plot based on Bray-Curtis distance, with colors for different datasets. The variance explained by populations (PERMANOVA *R*^2^) and its significance (PERMANOVA *p* value) were annotated in the figure. (**b**) Average Bray-Curtis distances between pairs of CRC datasets. Values on the diagonal referred to average Bray-Curtis distances between samples within the same dataset. Off-diagonal values refer to average Bray-Curtis distances between pairs of samples in different datasets. Larger values indicated a more dispersed distribution (on-diagonal) or bigger differences (off-diagonal).

To investigate the impact of differences between two populations on cross-study prediction, we create pseudo-population vectors *v*_*k*_, *k* = 1, 2:

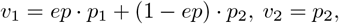

where *ep* is the population effect quantifying differences between two populations. Note that *v*_1_*− v*_2_ = *ep*·(*p*_1_*− p*_2_). Therefore, the differences between the two simulated populations increase with *ep*. At *ep* = 0, the two simulated populations share the same underlying distribution, resulting in no population differences between the training and testing datasets. Conversely, at *ep* = 1, the simulated populations exhibit the largest possible differences. In our simulations, we examined the overall trend for different normalization methods by varying *ep* from 0 to 1 in increments of 0.2. For scaling methods and transformation methods that work effectively at smaller *ep* values, we set *ep* to range from 0 to 0.25 in increments of 0.05.

Out of the 154 shared taxa between the two populations, we randomly selected 10 taxa and hypothesized that these taxa were associated with a specific disease of interest. Considering that disease-associated taxa can either be enriched or depleted, we presumed the first 5 taxa to be enriched and the latter 5 to be depleted. These 10 taxa were fixed in the following analysis. The abundance vectors for simulated controls of selected disease-associated taxa were not changed (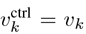, *k* = 1, 2), while the abundance vectors for simulated cases of selected disease-associated taxa were defined as follows:

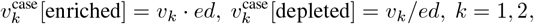

where *ed ∈ {*1.02, 1.04, 1.06*}* denoted a disease effect factor that quantified the differences between cases and controls. As the value of *ed* increases, the difference between case and control samples becomes more marked. Once we had the new vectors, we re-normalized them into probability vectors denoted as 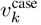, *k* = 1, 2.

Pseudo probability for control sample *j* in population *k*, denoted as 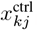, was generated under the assumption of a Dirichlet distribution: 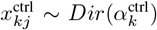, with 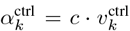 for *k* = 1, 2. When *c* is very large, the variance of 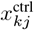 will be close to 0, and it is similar to 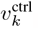. To introduce some variability while generating non-zero probabilities, we set *c* to 1*×* 10^6^. The read counts for control sample *j* in population *k* was subsequently simulated using multinomial distribution, with a library size of 1, 000, 000, described by:

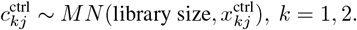

The generation of case samples followed a similar procedure, with the creation of 50 control and 50 case samples within each population.

### Scenario 2: Different batch effects in studies with the same background distribution of taxa in populations

In this scenario, we utilized Feng dataset [25] as the template for simulations. This ensured that the background distribution remained consistent between the training and testing datasets, thereby eliminating the population effects discussed in Scenario 1. We generated the read counts of training and testing data with 50 controls and 50 cases each by following the same procedure described in Scenario 1. It involved using multinomial distributions with a sample size of one million reads. The number of disease-associated taxa was set to 10 and disease effects varied from 1.02 to 1.06 with increments of 0.2.

To simulate batch effects, we followed a similar procedure as in Zhang et al [69]. They used the linear model assumed in the ComBat batch correction method [61] as the data-generating model for batch effects. Specifically, we assumed that both the mean (*γ*_*ik*_) and variance (*δ*_*ik*_) of taxon *i* were influenced by the batch *k*. The values of *γ*_*ik*_ and *δ*_*ik*_ were randomly drawn from normal and inverse gamma distributions:

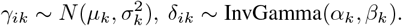

To set the hyper-parameters (*μ*_*k*_, *σ*_*k*_, *α*_*k*_, *β*_*k*_), we specify two values to represent the severity of batch effects. This included three levels for batch effects on the mean (*sev*_*mean*_ *∈ {*0, 500, 1000 *}*) and three levels for batch effects on the variance (*sev*_*var*_ *∈ {*1, 2, 4 *}*). For each severity level, the variance of *γ*_*ik*_ and *δ*_*ik*_ was fixed at 0.01. The parameters are then added or multiplied to the expression mean and variance of the original study. The batch effects were only simulated on the training data while the test dataset was unchanged.

### Scenario 3: Different disease models of studies with the same background distribution of taxa in populations

In this scenario, we hypothesized that the model for disease-associated taxa could vary between populations. To avoid the population effects described in Scenario 1, we utilized the Feng dataset [25] as template for simulations. To avoid the batch effects described in Scenario 2, no batch effects were introduced into this simulation scenario.

For the selection of disease-associated taxa, we predefined 10 taxa for the training data. A subset of taxa was chosen from the initially selected 10 and additional taxa were included to maintain a total of 10 signature taxa in the testing data. The degree of similarity between the training and testing data was determined by the number of overlapping taxa, ranging from 2 to 10 with increments of 2. Subsequently, the two populations were simulated following the same procedure as in the previous two scenarios. The simulation parameters included 100 samples per population (50 controls and 50 cases), one million reads per sample, and a disease effect of 1.02, 1.04, 1.06.

## Results

### Different datasets have different background distributions

There are eight publicly accessible colorectal cancer (CRC) datasets shown in Table 1, including Feng [25], Gupta [26, 68], Thomas [8], Vogtmann [28], Wirbel [29], Yachida [30], Yu [9], and Zeller [31]. In total, we included 1260 samples (625 controls, 635 CRC cases) from multiple countries such as the USA, China, France, etc. The participant demographics ranged from 21 to 90 years, with a male representation of 59.6%. The datasets were characterized by diverse body mass index (BMI) values and included subjects with other health conditions such as hypertension, hypercholesterolemia, and Type 2 Diabetes (T2D). DNA extraction and sequencing were conducted using various protocols and platforms. Our analysis aimed to examine the background distribution differences among these datasets.

In order to assess population differences across the CRC datasets, a PCoA plot based on Bray Curtis distance was generated. Figure 1(**a**) revealed distinct separations between different datasets, suggesting variations in microbial composition among the populations. Although the observed separation accounted for a small proportion (7.9%) of the total variance, statistical significance was confirmed through the PERMANOVA test (*p* = 0.001). These findings underscored the substantial heterogeneity in microbial communities across diverse CRC datasets, despite the relatively modest contribution to the overall variance. To quantify the overlaps of these datasets, we computed the average Bray-Curtis distance (Figure 1(**b**)). The dispersion of individual datasets was represented on the diagonal, with the largest dispersion observed in the Gupta dataset. Among the off-diagonal values that measured the average distance between samples in different datasets, Feng and Gupta exhibited the lowest overlap, with a distance of 0.901. Consequently, controls from these two datasets were selected as the template data for subsequent simulations in scenario 1. Mixing these two populations with decided proportions allowed us to control the heterogeneities between simulated populations.

Our analysis also extended to five distinct IBD datasets, as depicted in supplementary Table S1. These included the Hall [32], HMP [70, 5], Ijaz [33], Nielsen [35], and Vila [6] datasets. Similar to the CRC datasets, the IBD datasets exhibited variations in geographical origin, age, BMI, and sequencing platforms. Supplementary Figure S1 revealed a clear separation between the different datasets (supplementary Figure S1(**a**)) along with evident dataset dispersion variations (supplementary Figure S1(**b**)). These observations underscore the fact that distinctive populations are inherently marked by their unique background distributions, a factor that must be judiciously accounted for in any microbiome-related analysis.

### Transformation and batch correction methods could enhance prediction performance for heterogeneous populations

In Scenario 1, the effects of different normalization methods on the prediction of binary phenotypes across diverse background distributions of taxa were investigated. The corresponding results are presented in Figure 2 with columns for population effects and panels for disease effects. When there were no population effects between the training and testing datasets (*ep* = 0), all normalization methods exhibited satisfactory performance, with average AUC values consistently achieving the maximum value of 1. However, as the population effects increased or disease effects decreased, an evident decline in predictive accuracy was observed.

**Figure 2:**
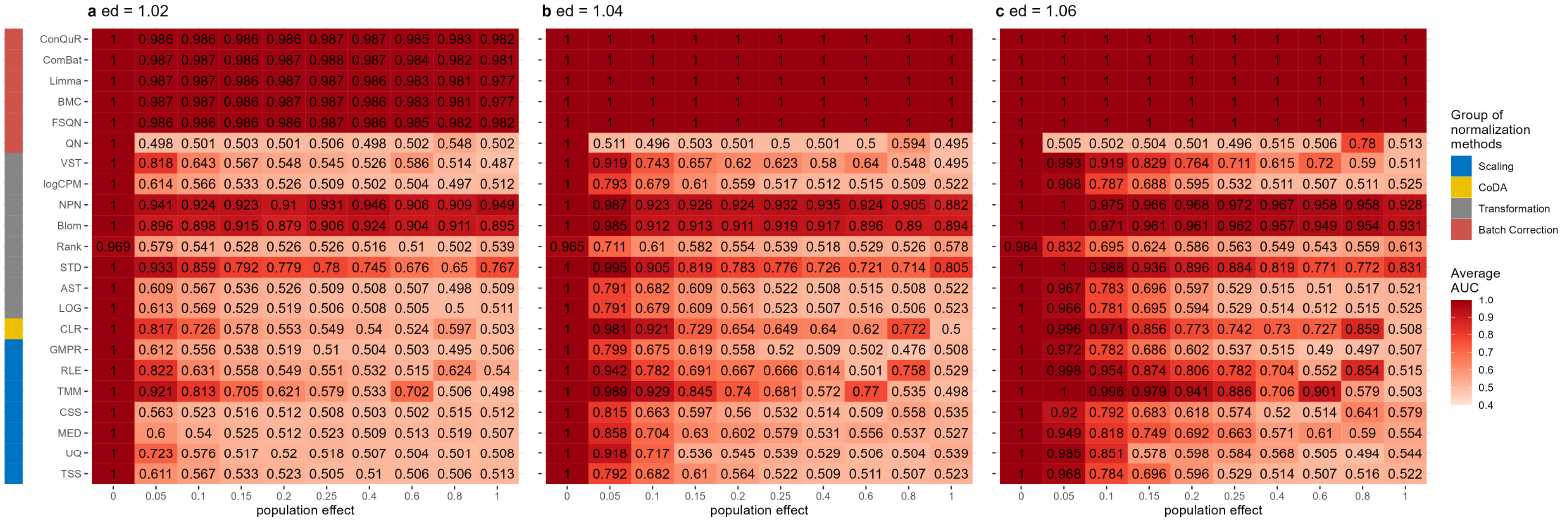
Heatmaps depicting average AUC values obtained from abundance profiles normalized by various methods for predicting simulated cases and controls in Scenario 1. The panels (**a**), (**b**), and (**c**) correspond to disease effects of 1.02, 1.04, and 1.06 respectively. The columns represent different values of population effects, while the rows represent different normalization methods, grouped based on their classifications in the left column.

When the differences between case and control were small (Figure 2(**a**), *ed* = 1.02), the prediction performance of scaling methods rapidly declined to 0.5 (random prediction value) as *ep* increased. TMM and RLE demonstrated better performances than TSS-based methods in a wider range of conditions. Notably, TMM maintained an AUC value above 0.6 when *ep <* 0.2. As disease effects increased (Figure 2(**b**) *ed* = 1.04 and (**c**) *ed* = 1.06), TMM and RLE exhibited superior ability to remove sample differences for predictions compared to TSS-based methods, indicating that normalization methods designed for RNA-Seq data distribution adjustment also benefited microbiome data predictions.

While normalized counts are commonly used for analyzing microbiome data, they still exhibit skewed distributions, unequal variances, and extreme values, which may limit their effectiveness in situations with significant heterogeneity. To enhance cross-population prediction performance, we applied various commonly used transformations, including CLR, LOG, AST, STD, Rank, Blom, NPN, logCPM, and VST. These transformation methods aimed to address one or several problems. For instance, logCPM and LOG transformations resolved skewness and extreme values, STD focused on unequal variances, VST tackled unequal variances and extreme values, and AST, CLR, Rank, Blom, and NPN addressed all three issues. The yellow and grey bars in Figure 2 represent the average prediction AUC values obtained using abundance profiles transformed by different methods. LOG, AST, Rank, and logCPM showed performances similar to TSS, indicating a failure in distribution adjustment. Conversely, transformation methods that achieved data normality, such as Blom and NPN, effectively aligned the data distributions across different populations for both population effects (*ep*) and disease effects (*ed*). Additionally, STD generally improved prediction accuracy, while the performance of CLR and VST transformation decreased with increasing population effects (*ep*).

Surprisingly, the batch correction methods denoted in red in Figure 2 yielded promising prediction results, with the exception of QN. QN forced the distribution of each sample to be the same, potentially distorting the true biological variation between case and control samples, making it difficult for the classifier to distinguish between the groups. While QN was only effective when the two populations originated from the same distribution, FSQN, BMC, limma, ComBat, and ConQuR significantly enhanced the reproducibility of response predictions, remaining unaffected by disease effects and population effects.

### Batch correction methods can successfully remove batch effects within the same population

In Scenario 2, we examined studies within the same population that exhibited technical variations and differences across batches. These batch effects can lead to substantial heterogeneity among the data batches [71]. Figure 3 showed the average AUC values obtained from random forest models using abundance profiles normalized by various methods across 100 runs. Overall, the prediction accuracy demonstrated an upward trend with increasing disease effects. However, the normalization methods exhibited varying responses to changes in batch means and variances.

**Figure 3:**
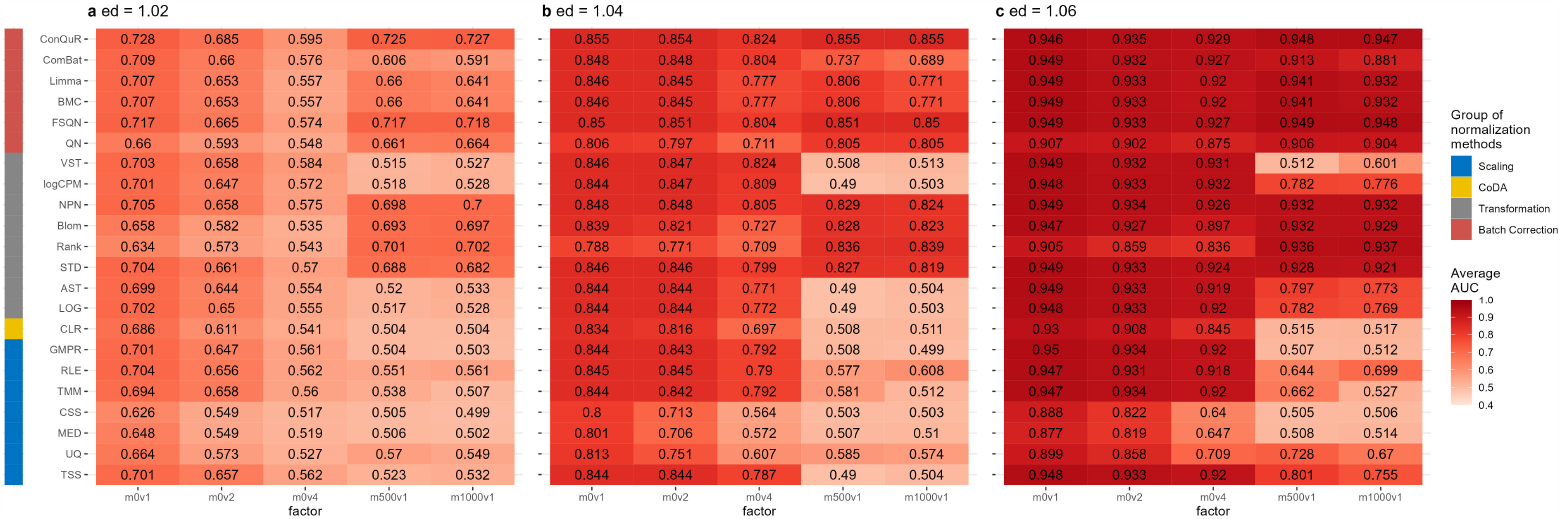
Heatmaps depicting average AUC values obtained from abundance profiles normalized by various methods for predicting simulated cases and controls in Scenario 2. The panels (**a**), (**b**), and (**c**) correspond to disease effects of 1.02, 1.04, and 1.06 respectively. The columns represent different combinations of batch mean and batch variation, with “m” for batch mean adjusting the mean and “v” for batch variance adjusting the variance. The rows represent different normalization methods, grouped based on their classifications in the left column.

Figure 3(**a**) displayed the results obtained with disease effect equal to 1.02. When the batch variance remained fixed (*sev*_*var*_ = 1), pronounced response to additive batch means (*sev*_*mean*_ = 0, 500, 1000) was observed among the scaling methods and some transformation methods (CLR, LOG, AST, logCPM, VST). These methods exhibited a decrease in AUC scores from approximately 0.7 to around 0.5 when *sev*_*mean*_ ≠ 0. In contrast, the STD, Rank, Blom, NPN, and all batch correction methods maintained a more robust level of AUC values (around 0.7) in the presence of varying batch means, as long as the batch variances did not change. These trends persisted with increasing disease effects, as depicted in Figure 3(**b**) and 3(**c**). Notably, among the methods more sensitive to batch means, scaling methods such as TMM and RLE exhibited a slight improvement in predictive accuracy as the batch means increased. Transformation methods like LOG, AST, and logCPM performed similarly.

The effects of batch variances on binary phenotype prediction remained consistent across different normalization methods. In Figure 3(**a**), when the batch mean was fixed at 0 and the batch variances were adjusted from 1 to 4, all normalization methods experienced an average decrease in AUC values of approximately 0.1. Among the scaling methods, namely MED, UQ, and CSS, which modified the scaling factor from TSS, consistently yielded lower AUC values compared to other methods for different batch variances (*sev*_*var*_ = 1, 2, 4). In Figure 3(**c**), with *ed* = 1.06, the influence of increased batch variance on prediction accuracy was reduced, indicating the dominance of disease effect in prediction. Most normalization methods achieved AUC scores above 0.9 when *sev*_*var*_ = 4, indicating successful removal of batch effects for predictions. Nonetheless, MED, UQ, and CSS continued to exhibit inferior ability in removing batch effects compared to other methods.

### The impact of disease model can be reduced by disease effects

In Scenario 3, we explored the influence of differences in disease models between the training and testing data on the prediction AUC scores. The results are presented in Figure 4. The overall trends in the relative performance of different normalization methods were consistent with the previous two scenarios. The AUC scores increased as the disease effects increased. And as expected, the AUC scores also increased as the number of overlapping disease-related taxa increased. For example, when *ed* = 1.02 (Figure 4(**a**)), the AUC values obtained using abundance profiles normalized by different methods were all approximately 0.6 when there were 2 overlapping disease-associated taxa between the training and testing data. When the number of disease-associated taxa increased to 10, the optimal AUC scores increased to 0.7. The same pattern was observed with *ed* = 1.04 and *ed* = 1.06. When the disease effects increased to 1.06 (Figure 4(**c**)), the majority of normalization methods achieved AUC scores exceeding 0.8, even when there were only 2 overlapped disease-associated taxa. This indicates that the impact of the disease model can be mitigated by stronger disease effects.

**Figure 4:**
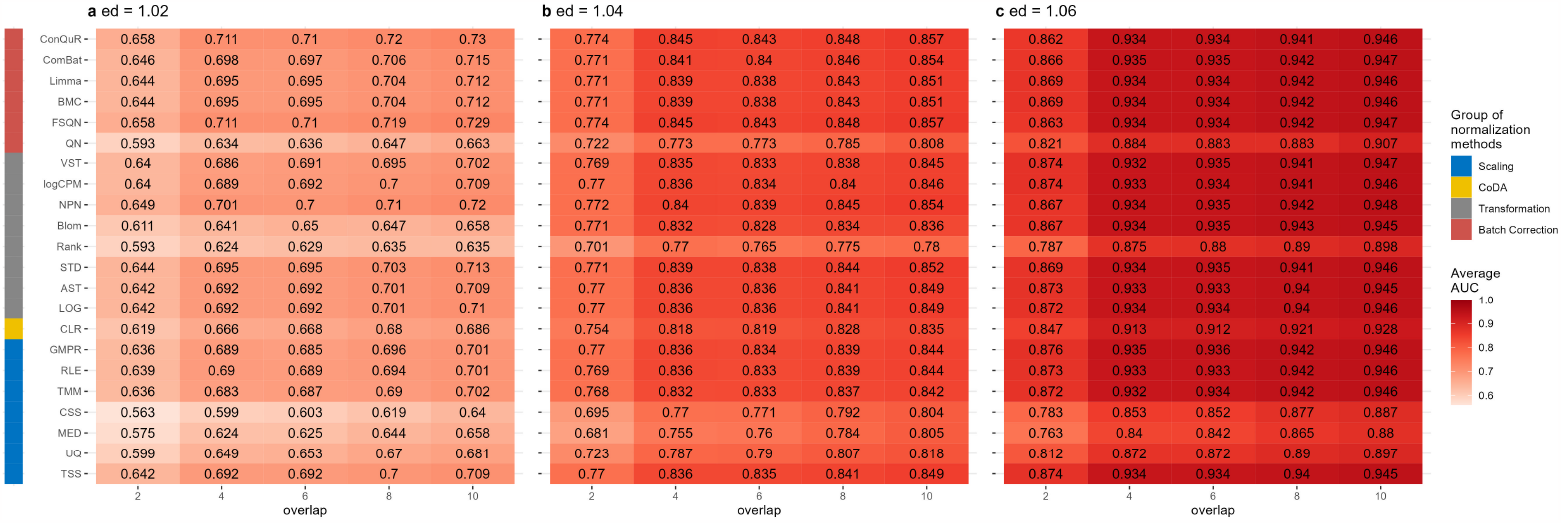
Heatmaps depicting average AUC values obtained from abundance profiles normalized by various methods for predicting simulated cases and controls in Scenario 3. The panels (**a**), (**b**), and (**c**) correspond to disease effects of 1.02, 1.04, and 1.06 respectively. The columns represent different numbers of overlapping disease-associated taxa in the training and testing datasets. The rows represent different normalization methods, grouped based on their classifications in the left column.

Figure 4 also illustrated that among the normalization methods we compared, scaling methods such as UQ, MED, and CSS had lower AUC values compared to other methods, as observed in the other two scenarios. QN also exhibited lower prediction performances. The other methods showed similar prediction performances with respect to different disease effects and different numbers of disease-associated taxa.

### Batch correction methods are necessary for cross-dataset predictions

We next evaluate various normalization methods using 8 gut microbiome datasets from shotgun sequencing related to CRC (Table 1). These experimental datasets were retrieved from the R package curatedMetagenomicData with a sample size larger than 30 for either cases or controls. Datasets were paired with one for model training and the other for validation. For each method, the AUC score based on the normalized abundance using random forest was calculated. We repeated the predictions 30 times to account for the randomness of the prediction model and the average of the AUC scores was reported for each study.

Supplementary Figure S2 presents box plots showing the AUC values obtained from the 30 repeated predictions. We observed unstable AUC values for most normalization methods when trained or tested on the Gupta dataset. This observation aligns with the data distribution depicted in Figure 1, where Gupta exhibited the greatest dissimilarities and variability compared to other datasets. The same observation holds true for the Feng dataset. Overall, none of the normalization methods consistently improved the prediction AUC values to a specific level. The prediction accuracy remained dependent on both biological and technical factors. For example, when the model was trained on Gupta and tested on Feng, most methods yielded average AUC scores around 0.7, except for Rank and VST (Figure 5(**a**)). None of the normalization methods achieved an AUC value above 0.8 to significantly improve prediction performance.

**Figure 5:**
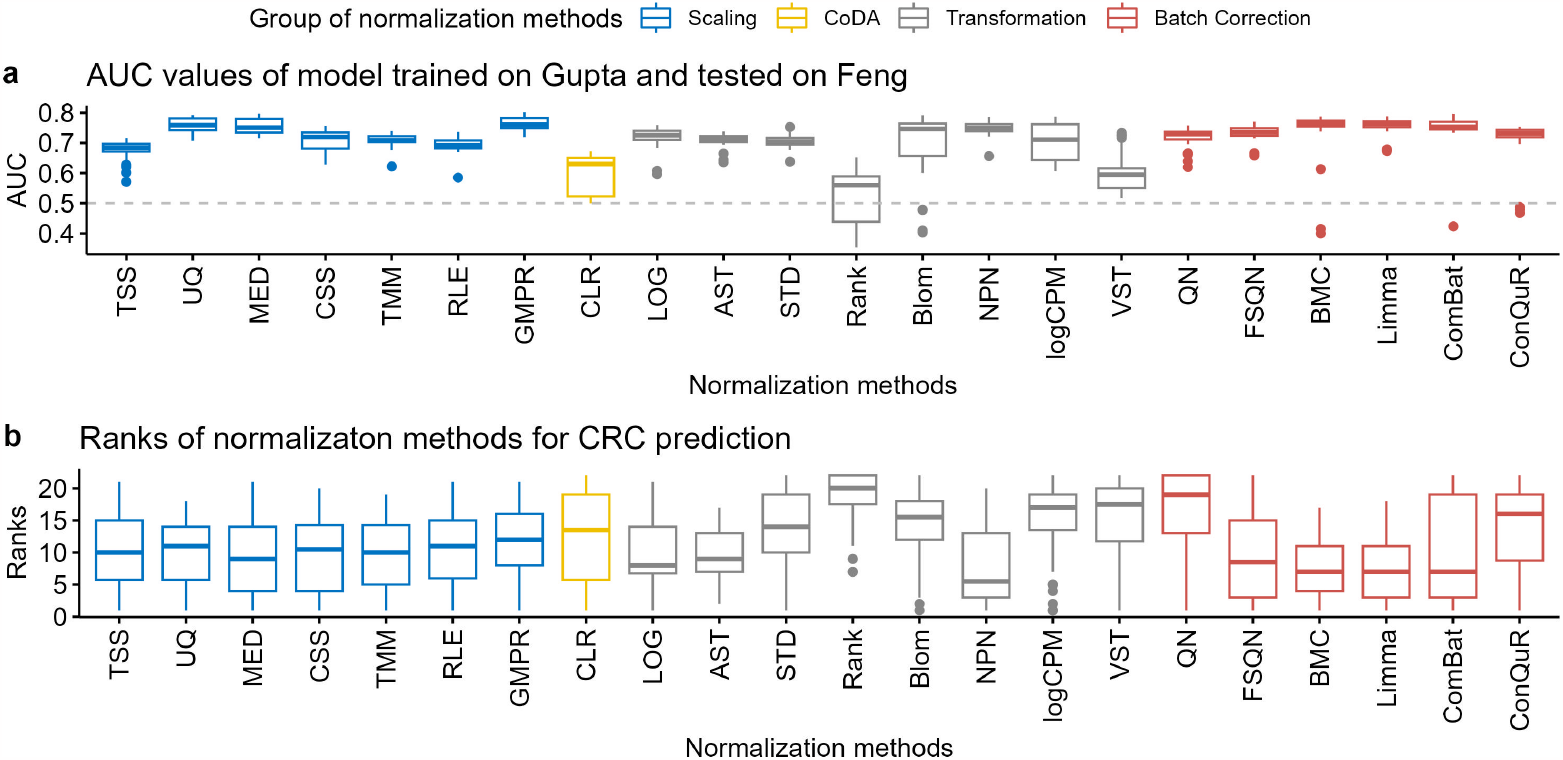
Comparison of normalization methods based on 8 CRC datasets in predictions. (**a**) Boxplots for AUC values of the model trained on Gutpa and tested on Feng. (**b**) Distribution of the ranks for different normalization methods according to the average AUC value under the same training and testing datasets.

To quantify the performance of normalization methods, we ranked all normalization methods based on the average AUC scores when the model was trained on the same dataset and validated on the same test dataset. The distributions of their ranks across these 8 studies for each method are depicted in Figure 5(**b**). A higher ranking (lower values in the box plot) indicates a better prediction performance. Among the twenty-two normalization methods we compared, batch correction methods, including FSQN, BMC, Limma, and ComBat, tended to have higher AUC values than other methods. However, the performance of ComBat displayed some fluctuations. Scaling methods ranked behind batch correction methods and performed similarly to each other in CRC dataset predictions, indicating relatively small population effects in CRC datasets.

We also applied the normalization methods to IBD datasets listed in supplementary Table S1 and conducted cross-dataset predictions. Supplementary Figure S3 displays the box plots of AUC values from the 30 repeated predictions, while supplementary Figure S4 shows the distributions of ranks for each method in pairs of IBD datasets. The results obtained were similar to those observed in the CRC dataset predictions. Among all the normalization methods, batch correction methods, including BMC and Limma, consistently demonstrated the best performance. Scaling methods, such as TMM and RLE, followed closely behind. However, FSQN and ComBat exhibited variable performance, occasionally achieving good results while sometimes yielding poor results. Overall, the trends in IBD dataset predictions were consistent with the observations made in CRC dataset predictions.

## Discussion

In our study, we conducted a comprehensive evaluation of various normalization methods for predicting binary phenotypes using both simulated and real microbiome data. The results revealed important insights into the performance and suitability of different normalization approaches in the context of disease prediction.

Our findings demonstrated that no single normalization method consistently outperformed others across all datasets and phenotypic outcomes. This suggests that the choice of normalization method should be carefully considered based on the specific dataset characteristics and research objectives. However, certain trends and patterns did emerge from our analysis.

Among the scaling methods, methods such as TMM and RLE performed comparably well, indicating their effectiveness in reducing technical variations and improving the comparability of data across samples. These methods are relatively simple and straightforward to implement, making them practical choices for normalization in microbiome data analysis.

Interestingly, compositional data analysis methods, CLR, exhibited mixed performance across different datasets. While it has been widely used in microbial community analysis, our results suggest that its effectiveness in disease prediction may vary depending on the specific dataset and phenotypic outcome. Further investigation is needed to understand the underlying factors influencing the performance of compositional data analysis methods in predicting binary phenotypes.

Transformation methods, including NPN and Blom, showed promising results in some datasets, highlighting their potential to improve prediction accuracy by capturing nonlinear relationships and addressing skewed distributions. These methods offer flexibility in handling diverse data types and can be particularly valuable in situations where data transformation is necessary to meet model assumptions.

Batch correction methods, such as BMC and Limma, consistently performed well across multiple datasets. These methods effectively accounted for batch effects, which are often present in multi-center or multicohort studies. The ability to remove batch effects is critical in ensuring accurate and reliable predictions, especially when integrating data from different sources.

It is worth noting that the performance of normalization methods was influenced by the heterogeneity of the datasets. In datasets where there were substantial biological and technical variations, the prediction accuracy remained primarily determined by these factors rather than the choice of normalization method. This emphasizes the importance of considering population effects, disease effects, and batch effects when interpreting the results of microbiome data analysis and developing predictive models.

Overall, our study underscores the need for careful consideration and evaluation of normalization methods in microbiome data analysis, particularly in the context of disease prediction. Researchers and practitioners should take into account the specific characteristics of their datasets, including population heterogeneity, disease effects, and technical variations when selecting and applying normalization methods. Additionally, future research should focus on developing novel normalization approaches that are tailored to the unique challenges of microbiome data and explore their performance in larger and more diverse datasets.

In conclusion, our comprehensive evaluation of normalization methods provides valuable insights into their performance in predicting binary phenotypes using microbiome data. This research contributes to the advancement of robust and reliable methodologies in microbiome research and paves the way for more accurate disease prediction and personalized therapeutic interventions based on the human microbiome.

## Data availability

All the CRC and IBD datasets used in this study are available in the R package curatedMetagenomicData (v3.8.0). All the codes used in the analysis can be found at https://github.com/wbb121/Norm-Methods-Comparison.

## Author contributions

F.S. and Y.L. designed and supervised the study. B.W. implemented the methods, conducted the computational analysis, and drafted the manuscripts. F.S. and Y.L. modified and finalized the manuscripts. All authors read and approved the final version of the manuscript.

## Additional information

