## Supplementary Table S1, Supplementary Figure S1, Supplementary Figure S2, Supplementary Figure S3, Supplementary Figure S4 for "Comparison of the effectiveness of different normalization methods for metagenomic cross-study phenotype prediction under heterogeneity"

### Supplementary Tables

Table S1: Characteristics of IBD datasets, with different DNA extraction kits (DNA-Exk) and sequencing platforms (Seq-Plat).

| Dataset | Country | No. of controls | No. of cases | DNA-Exk | Seq-Plat | Reference |
|---|---|---|---|---|---|---|
| Hall | United States of America | 74 | 185 | other/Qiagen | IlluminaMiSeq | [1] |
| HMP | United States of America | 426 | 1201 | Chemagen | IlluminaHiSeq | [2, 3] |
| Ijaz | United Kingdom | 38 | 56 | NA | IlluminaHiSeq | [4] |
| Nielsen | Denmark/Spain | 248 | 148 | NA | IlluminaHiSeq | [5] |
| Vila | Netherlands | 1135 | 355 | Qiagen | IlluminaHiSeq | [6] |

### Supplementary Figures

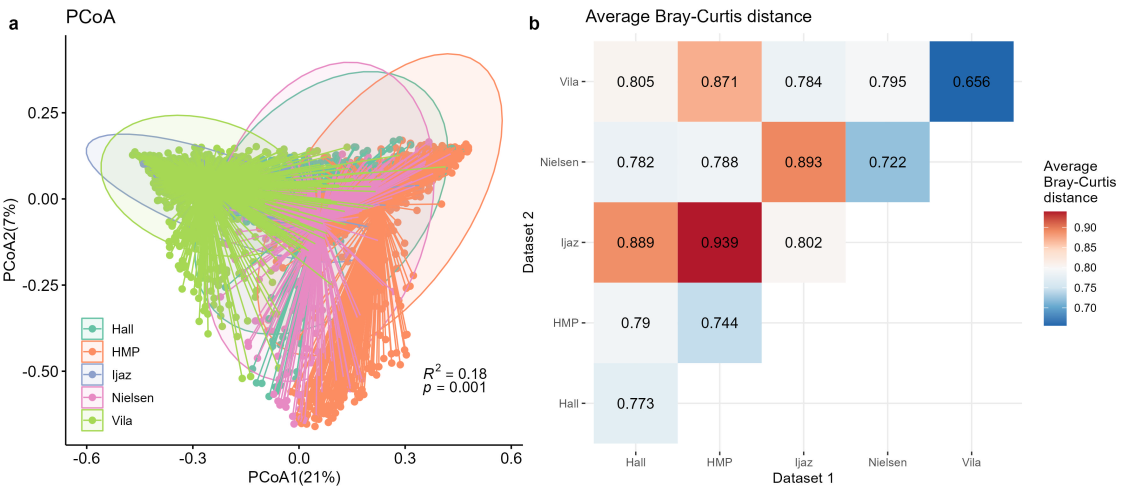


Figure S1: Different IBD populations had different background distribution patterns. (**a**) PCoA plot based on Bray-Curtis distance, with colors for different datasets. The variance explained by populations (PERMANOVA $R^2$) and its significance (PERMANOVA $p$ value) were annotated in the figure. (**b**) Average Bray-Curtis distances between pairs of IBD datasets. Values on the diagonal referred to average Bray-Curtis distances between samples within the same dataset. Off-diagonal values refer to average Bray-Curtis distances between pairs of samples in different datasets. Larger values indicated a more dispersed distribution (on-diagonal) or bigger differences (off-diagonal).

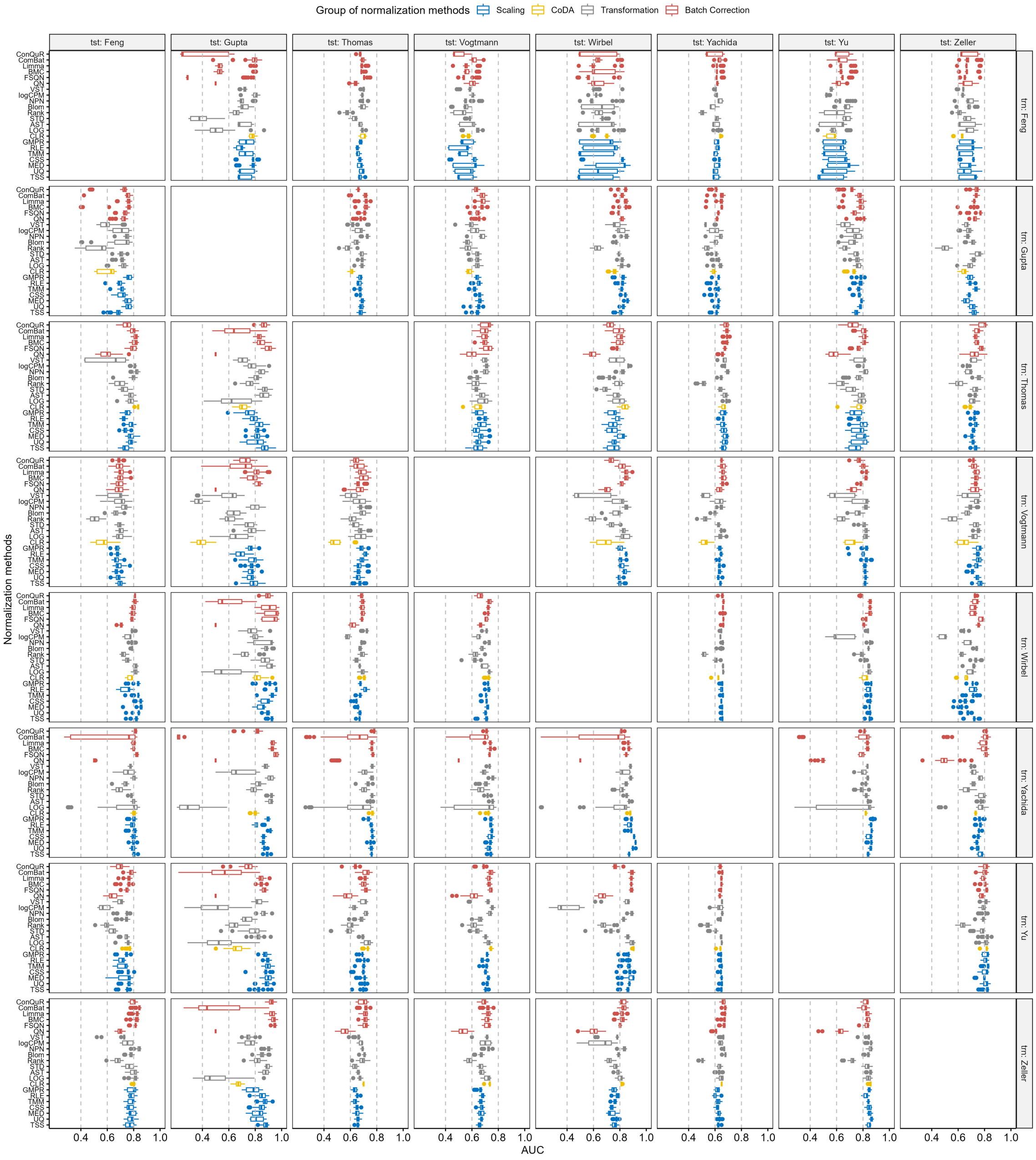


Figure S2: Box plots for cross-dataset predictions of disease status using abundance profiles normalized by various methods in CRC datasets, with panel rows representing training data and panel columns representing testing data. The normalization methods were categorized and color-coded by their respective groups.

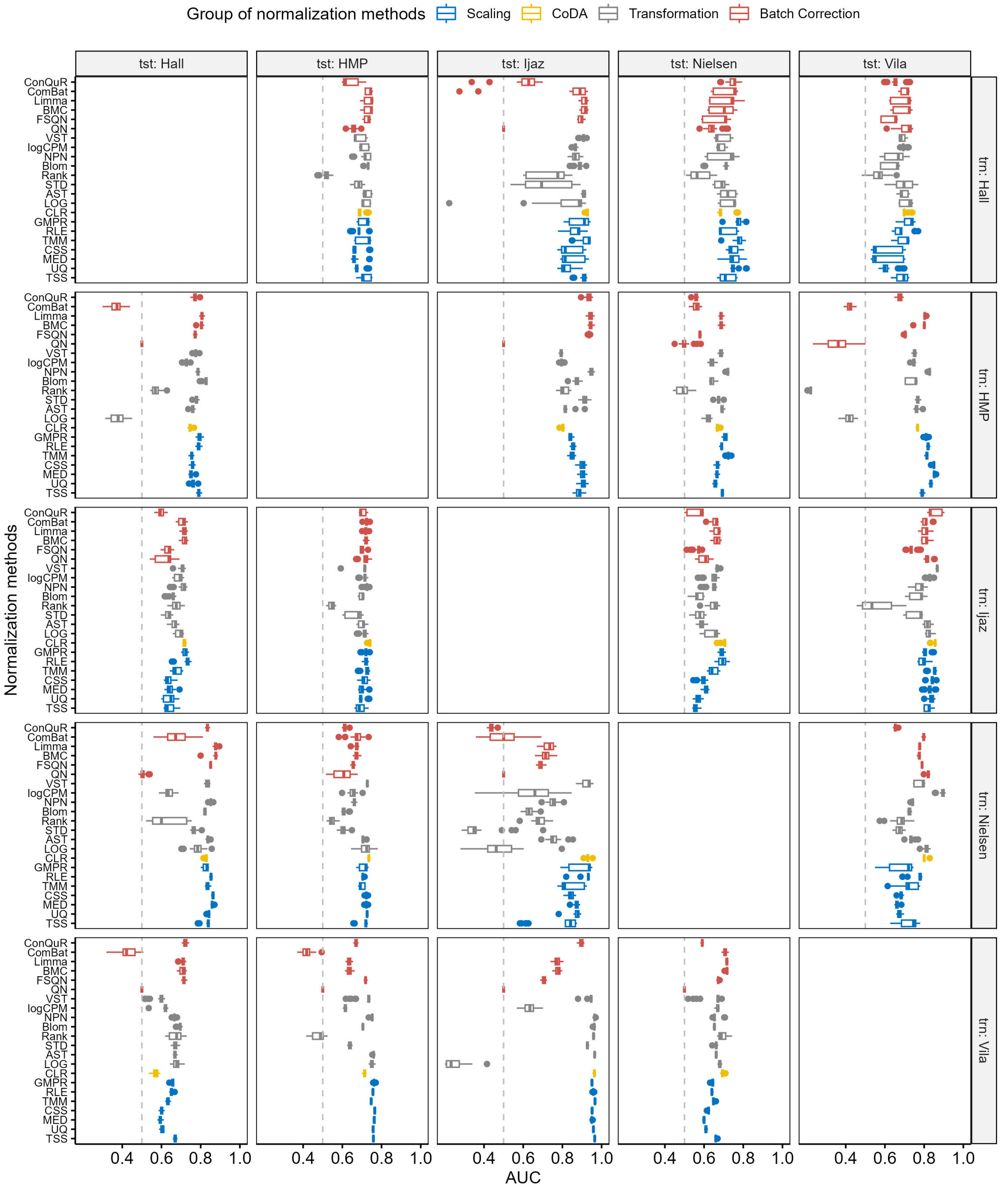


Figure S3: Box plots for cross-dataset predictions of disease status using abundance profiles normalized by various methods in IBD datasets, with panel rows representing training data and panel columns representing testing data. The normalization methods were categorized and color-coded by their respective groups.

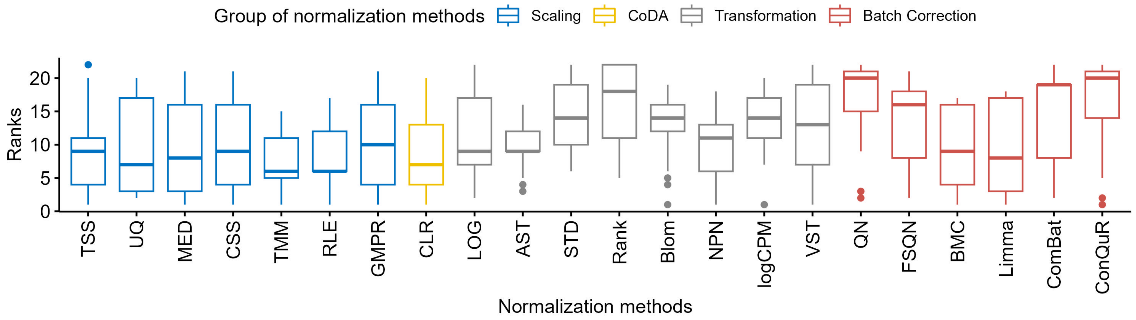


Figure S4: Distribution of the ranks for different normalization methods according to the average AUC value under the same training and testing datasets of IBD datasets. The average AUC values are ranked in descending sort with rank 1 for the highest average AUC value.
